# A large-scale species level dated angiosperm phylogeny for evolutionary and ecological analyses

**DOI:** 10.1101/809921

**Authors:** Steven B. Janssens, Thomas L.P. Couvreur, Arne Mertens, Gilles Dauby, Samuel Vanden Abeele, Filip Vandelook, Hans Beeckman, Maurizio Mascarello, Vincent Droissart, Marc S.M. Sosef, Michelle van der Bank, Olivier Maurin, William Hawthorne, Cecily Marshall, Maxime Réjou-Méchain, Denis Beina, Fidele Baya, Léo-Paul M.J. Dagallier, Vincent S.F.T. Merckx, Brecht Verstraete, Olivier Hardy

**Affiliations:** Meise Botanic Garden, Nieuwelaan 38, BE-1860 Meise, Belgium; Plant Conservation and Population Biology, KU Leuven, Kasteelpark Arenberg 31, PO Box 2435, 3001 Leuven, Belgium; IRD, DIADE, University of Montpellier, Montpellier, France; UMR AMAP, IRD, F-34000 Montpellier, France; Royal Museum for Central Africa, 3080 Tervuren, Belgium; Africa & Madagascar Department, Missouri Botanical Garden, St Louis, MO, USA; Herbarium et Bibliothéque de Botanique Africaine, Université Libre de Bruxelles, Bruxelles, Belgium; Department of Botany and Plant Biotechnology, University of Johannesburg, Auckland park, Johannesburg PO Box 524, South Africa; Department of Comparative Plant and Fungal Biology, Royal Botanic Gardens, Kew, Richmond, Surrey TW9 3AE, UK; Department of Plant Sciences, University of Oxford, South Parks Road, Oxford OX1 3RB, UK; UPR BSEF, CIRAD, Campus International de Baillarguet, F-34398 Montpellier, France; Université de Bangui – Cerphameta, BP 1450 Bangui, Central African Republic; Ministère des Eaux, Forêts, Chasse et Pêche, BP 3314 Bangui, Central African Republic; Naturalis Biodiversity Center, Leiden University, PO Box 9517, 2300RA Leiden, the Netherlands; Natural History Museum, University of Oslo, Oslo, Norway; Laboratoire d’Evolution Biologique et Ecologie, Faculté des Sciences, Université Libre de Bruxelles, Brussels, Belgium

**Keywords:** phylogeny, angiosperms, large-scale dating analyses, evolution, ecology

## Abstract

Phylogenies are a central and indispensable tool for evolutionary and ecological research. Even though most angiosperm families are well investigated from a phylogenetic point of view, there are far less possibilities to carry out large-scale meta-analyses at order level or higher. Here, we reconstructed a large-scale dated phylogeny including nearly 1/8th of all angiosperm species based on two plastid barcoding genes, *matK* and *rbcL.* Novel sequences were generated for several species, while the rest of the data were mined from GenBank. The resulting tree was dated using 56 angiosperm fossils as calibration points. The resulting megaphylogeny is one of the largest dated phylogenetic tree of angiosperms yet, consisting of 36,101 sampled species, representing 8,399 genera, 426 families and all orders. This novel framework will be useful to investigate different broad scale research questions in ecological and evolutionary biology.

## Introduction

During the past two decades, awareness has grown that ecological and evolutionary studies benefit from incorporating phylogenetic information (*Wanntorp et al. 1990, Webb et al. 2002*). In some ecological disciplines it has even become almost unimaginable that a spatiotemporal context is not considered when specific hypotheses are tested. For example, in the fields of community ecology, trait-based ecology and macroecology, macroevolutionary and historical biogeography research hypotheses cannot be properly tested without the incorporation of a phylogenetic framework (e.g. *Graham and Fine 2008, Hardy 2008, Kissling 2017, Vandelook et al. 2012, Vandelook et al. 2018, Couvreur et al. 2011, Janssens et al. 2009,Janssens et al. 2016*). Likewise, phylogenetic diversity is considered an important element in conservation biology and related biodiversity assessment studies (*Chave et al. 2007*). Even though the importance of phylogenetics in ecology and evolution is recognized, it remains somehow strenuous to combine ecological research with evolutionary biology and integrate it in a phylogenetic scenario. This discrepancy is sometimes caused by a lack of awareness and knowledge about the other disciplines, whereby researchers could be reluctant to reach out to such expertise and combine their results into new disciplines.. Also differences in methodologies and techniques applied by ecologists and evolutionary biologists can sometimes cause a certain hesitation to go for a complementary approach with blending disciplines. In addition, there is a nearly continuous development of new insights and techniques in the field of ecology and evolution (e.g. *Bouckaert et al. 2019, Revell et al. 2008, Revell 2012, Suchard et al. 2018*), making it rather challenging to keep up to date with the latest novelties. Furthermore, not all organisms investigated from an ecological perspective are present in molecular databases, which make it difficult to construct a perfectly matching phylogenetic hypothesis for further analysis. For scientists that focus on resolving specific evolutionary or ecological queries, building a phylogenetic framework from novel gene sequence data is often a heavy burden as it takes a lot of time, money, and effort, apart even from the specific expertise needed. The construction of a purpose-built phylogeny can be considered as rather costly and labour-intensive, and requires more elaborate expertise on novel techniques than when sequences are merely mined from GenBank in order to make a tree based on already existing sequences. Whereas the former strategy allows the user to make a tailor-made phylogeny that can be used for further ecological or evolutionary purposes, the latter is less proficient as one can only use the sequences that are available in genetic databases. Nevertheless, in case of large-scale meta-analyses becomes almost impossible to obtain sequence data from all species investigated. When there is a need to examine evolutionary and ecological trends in an historical context, a large-scale phylogenetic hypothesis that is optimized in a spatiotemporal context provides an optimal solution.

There is currently an ongoing quest to optimize the methodology for constructing large-scale mega-phylogenies that can be used for further ecological and evolutionary studies. This is done by either mining and analysing publicly available DNA sequences (*Zanne et al. 2014*), amalgamating published phylograms *Hinchliff et al. 2015*) or the combination of both (*Smith and Brown 2018*). For example, *Zanne et al. (2014)* constructed their own large supermatrix-based phylogeny that was used to gain more insights into the evolution of cold-tolerant angiosperm lineages. However, the study of *Qian and Jin (2016)* showed that the phylogeny of *Zanne et al. (2014)* contained several taxonomic errors. Also the approaches of *Smith and Brown (2018)* and *Hinchliff et al. (2015)* also do not always provide the most optimal phylogenetic framework for further analyses as both studies use a (partially) synthetic approach based on already published phylograms that can putatively contain inconsistencies in their estimated node ages. The main goal of the current study is therefore to provide a large-scale dated phylogeny - encompassing at least 1/10th of all angiosperms - that can be used for further ecological and evolutionary analyses. In order to construct this angiosperm phylogeny, a comprehensive approach was applied in which sequence data were both mined and generated, subsequently aligned, phylogenetically analysed and dated using over 50 fossil calibration points. With the applied methodology we aimed to create sufficient overlap in molecular markers without having too much missing sequence data in the datamatrix. In addition, phylogenetic analyses as well as the age estimation assessment were performed as a single analysis on the whole datamatrix in order to create a dated angiosperm mega-phylogeny that is characterized by a low degree of syntheticness.

## Material and Methods

### Marker choice

In 2009, the Consortium for the Barcode of Life working group (CBOL) advised sequencing of the two plastid markers *matK* and *rbcL* for identifying plant species, resulting in a massive amount of data available on GenBank. *rbcL* is a conservative locus with low level of variation across flowering plants and therefore useful to reconstruct higher level divergence. In contrast, *matK* contains rapidly evolving regions that are useful to study interspecific divergence (*Hilu et al. 2003, Kress et al. 2005*). Thus, the combination of *matK* and *rbcL* has the advantage of combining different evolutionary rates, making it possible to infer relationships at different taxonomic levels. In addition, we sampled only *matK* and *rbcL* markers in order to reduce missing data to a minimum due to its major influence on the phylogenetic relationships between species. These supermatrix approaches - which generally contain a substantial amount of missing data – can suffer from imbalance in presence/absence for each taxon per locus resulting in low resolution and support or even wrongly inferred relationships (*Sanderson and Shaffer 2002, Roure et al. 2013*).

### Taxon sampling

We extracted angiosperm sequence data of *rbcL* and *matK* from GenBank (February 15, 2015) using the ‘NCBI Nucleotide extraction’ tool in Geneious v11.0 (Auckland, New Zealand). Five gymnosperm genera were chosen as outgroup (*Suppl. material 1*). This large dataset was supplemented with 468 specimens of African tree species obtained via multiple barcoding projects (available at the Barcode of Life Data Systems (BOLD)) as well as via additional lab work (see paragraph on molecular protocols below). Newly obtained sequences are submitted to GenBank (*Suppl. material 1*).

### Molecular protocols

A modified CTAB protocol was used for total genomic DNA isolation (*Tel-Zur et al. 1999*). Secondary metabolites were removed by washing ground leaf material with extraction buffer (100mM TrisHCl pH 8, 5mM EDTA pH 8, 0.35M sorbitol). After the addition of 575µl CTAB lysis buffer with addition of 3% PVP-40, the samples were incubated for 1.5 hours (60°C). Chloroform-isoamylalcohol (24/1 v/v) extraction was done twice, followed by an ethanol-salt precipitation (absolute ethanol, sodium acetate 3M). After centrifugation, the pellet was washed twice (70% ethanol), air-dried, and dissolved in 100µl TE buffer (10mM TrisHCl pH 8, 1mM EDTA pH 8).

Amplification reactions of *matK* and *rbcL* were carried out using standard PCR (25µl). Reactions commenced with a 3 minute heating at 95°C followed by 30 cycles consisting of 95°C denaturation for 30s, primer annealing for 60s, and extension at 72°C for 60s. Reactions ended with a 3 minute incubation at 72°C. Annealing temperatures for *matK* and *rbcL* were set at 50°C and 55°C respectively. Primers designed by Kim J. (unpublished) were used to sequence *matK*, whereas *rbcL* primers were adopted from *Fay et al. (1997)* and *Little and Barrington (2003)*. PCR products are cleaned using an ExoSap purification protocol. Purified amplification products were sequenced by the Macrogen sequencing facilities (Macrogen, Seoul, South Korea). Raw sequences were assembled using Geneious v11.0 (Biomatters, New Zealand).

### Sequence alignment and phylogenetic analyses

We are aware that the publicly available databaseGenBank contains a large amount of erroneous data (*Ashelford et al. 2005, Yao et al. 2004, Shen et al. 2013*). Retrieving the sequence data was therefore subjected to a quality control procedure. All downloaded sequences were blasted (Megablast option) against the GenBank database, thereby discarding all sequences with anomalies against their original identification. Minimum similarity in BLAST was set at 0.0005, whereas word size (W) was reduced to 8 for greater sensitivity of the local pairwise alignment and the maximum hits was set at 250. A single sequence of each fragment was retained for each taxon name or non-canonical NCBI taxon identifier given in GenBank. Furthermore, sequences with multiple ambiguities were discarded as well as sequences with similar taxon names but different nucleotide sequences. In addition, sequences with erroneous taxonomic names (checked in R using the “Taxize” and “Taxonstand” packages (*R Development Core Team 2009, Cayuela and Oksanen 2016, Chamberlain et al. 2016*)) were removed from further analyses. Interestingly, Taxize uses the Taxonomic Name Resolution Service (TNRS) function to match taxonomic names, whereas Taxonstand is linked with ‘The Plant List’ database. As such we also checked the validity of the taxonomic names in our dataset using both databases. Only those taxa that have names that are considered valid for both databases were kept for further analyses.

For sequence fragments that are protein-encoded (complete *rbcL* and parts of *matK*), comparison of amino acid (AA) sequences based on the associated triplet codons between taxa was applied. As a result, taxa with a sudden shift in AA or frame shift were discarded from the dataset.

Alignment was carried out in multiple stages. Due to our large angiosperm-wide dataset, an initial alignment (automatically and manually) was conducted for each order included in the dataset. Subsequently, the different alignments were combined using the Profile alignment algorithm (Geneious v11.0, Auckland, New Zealand). The initial automatic alignment was conducted with MAFFT (*Katoh et al. 2002*) using an E-INS-i algorithm, a 100PAM/k=2 scoring matrix, a gap open penalty of 1.3, and an offset value of 0.123. Manual fine-tuning of the aligned dataset was performed in Geneious v11.0 (Auckland, New Zealand). During the manual alignment of the different datasets, we carefully assessed the homology of every nucleotide at each position in the alignment (*Phillips et al. 2009*). The large amount of angiosperm taxa included in the analyses often provided a good view on the evolution of the nucleotides at certain positions in which some taxa functioned as transition lineages between differing nucleotides and their exact position in the alignment. The importance of a well-designed homology assessment for a complex sequence dataset has been proven successful here for the phylogenetic inference of the angiosperms.

The best-fit nucleotide substitution model for both *rbcL* and *matK* was selected using jModelTest 2.1.4. (*Posada 2008*) out of 88 possible models under the Akaike information criterion (AIC). The GTR+G model was determined as the best substitution model for each locus. Maximum likelihood (ML) tree inference was conducted using the Randomized Axelerated Maximum Likelihood (RAxML) software version 7.4.2 (*Stamatakis 2006*) under the general time reversible (GTR) substitution model with gamma rate heterogeneity and lewis correction. Although the phylogeny based on the plastid dataset generated relationships that corresponded well with currently known angiosperm phylogenies (e.g. *Wikström et al. 2001, Soltis et al. 2002, Moore et al. 2007, Magallón and Castillo 2009, Magallón 2014, Magallón et al. 2015, Bell et al. 2005, Bell et al. 2010*), we decided to use a constraint in order to make sure that possible unrecognized mismatches for certain puzzling lineages were significantly reduced. The constraint tree follows the phylogenetic framework of APG4 (*Angiosperm Phylogeny Group 2016*) at order level. At lower phylogenetic level, families were only constrained as polytomy in their specific angiosperm order. Genera and species were not constrained.

Support values for the large angiosperm dataset were obtained via the rapid bootstrapping algorithm as implemented in RAxML 7.4.2 (*Stamatakis 2006*), examining 1000 pseudo-replicates under the same parameters as for the heuristic ML analyses. Bootstrap values were visualised using the Consensus Tree Builder algorithm as implemented in Geneious v11.0.

### Divergence time analysis

Evaluation of fossil calibration points was carried out following the specimen-based approach for assessing paleontological data by *Parham et al. (2012)*. As such, 56 angiosperm fossils were used as calibrations points in our molecular dating analysis. Detailed information about the fossils, including (1) citation of museum specimens, (2) locality and stratigraphy of fossils, (3) referenced stratigraphic age, and (4) crown/stem node position is provided in *Table 1*. Fossils are placed at both early and recently diversified lineages within the angiosperms. Due to the large size of the dataset, we applied the penalized likelihood algorithm as implemented in treePL (*Smith and O’Meara 2012*), which utilizes hard minimum and maximum age constraints. In order to estimate these hard minimum and maximum age constraints, we calculated the log normal distribution of each fossil calibration point using BEAUti v.1.10 (*Suchard et al. 2018*). Maximum age constraints for each fossil correspond to the 95.0% upper boundary of the computed log normal distribution in which the offset equals the age of the fossil calibration point, the mean is set at 1.0, and the standard deviation at 1.0. This methodology resulted in a minimum 15 million year broad interval for each angiosperm calibration point (*Table 1*). Due to recently published studies in which both old and young age estimates were retrieved for the crown node of the angiosperms (e.g. *Bell et al. 2005, Bell et al. 2010, Magallón et al. 2015, Magallón 2014, Magallón and Castillo 2009, Moore et al. 2007, Smith et al. 2010, Wikström et al. 2001, Soltis et al. 2002*), we opted to set the hard maximum and minimum calibration of the angiosperms at 220 and 180 million years, respectively. As for the overall calibration, we followed the strategy of *Smith et al. (2010)* in which all fossils were considered as a minimum-age constraint. *Smith et al. (2010)* applied this approach since earlier studies on angiosperm evolution had treated tricolpate fossil pollen as maximum-age constraint, thereby maybe artificially pushing the root age of the angiosperms towards more recent times (e.g. *Soltis et al. 2002, Magallón et al. 2015, Magallón 2014, Magallón and Castillo 2009, Moore et al. 2007, Bell et al. 2010, Bell et al. 2005*).

**Table 1.**
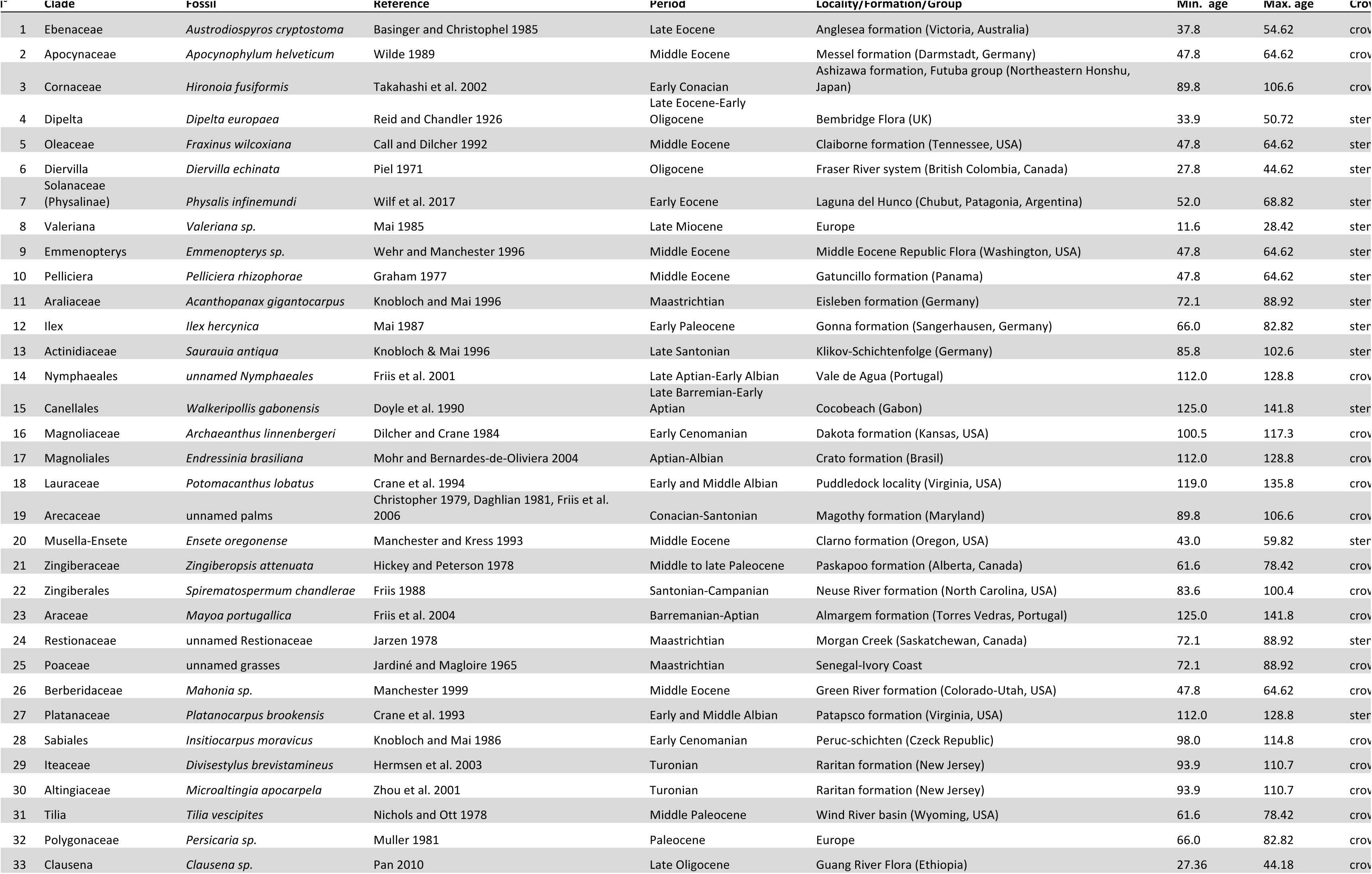
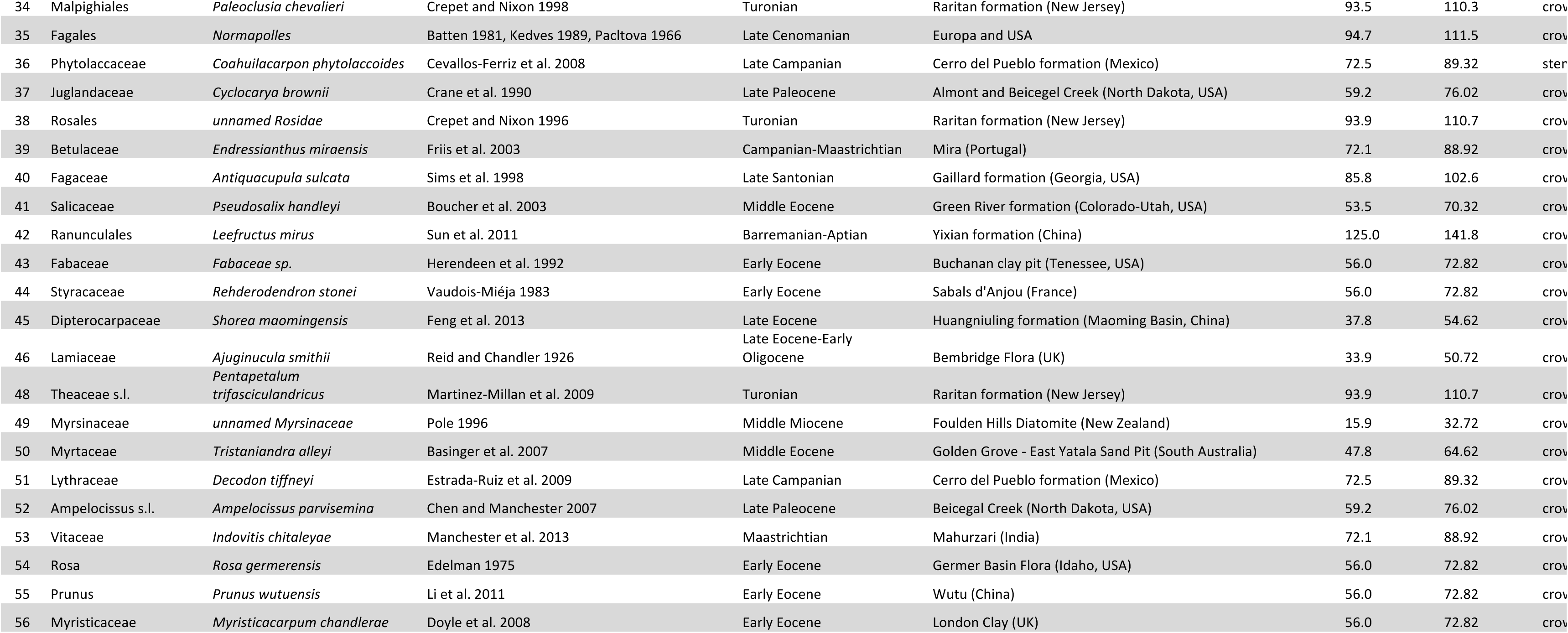
List of fossils used as calibration points, including their oldest stratigraphic occurrence, minimum and maximum ages, the calibrated clades and used references. cr.=crown, st.=stem.

## Results and Discussion

The final aligned data matrix consists of 36,101 angiosperm species. *matK* sequences were mined for 31,391 species (87%), whereas *rbcL* sequences were obtained for 26,811 (74%) species (*Suppl. material 1*). The sequence dataset has an aligned length of 4,968 basepairs (bp) of which 4,285 (86%) belong to *matK* and 683 (14%) to *rbcL*. Within *rbcL* all characters were variable (100%), whereas for *matK* 3,921 characters (91.5%) were variable. Support value analyses indicate that approximately 26% of the branches have a bootstrap value >75 (*Suppl. material 2*). Based on the different studies that estimated the total number of flowering plants currently described (between 260,000 and 450,000 species) (*Crane et al. 1995, Christenhusz and Byng 2016, Cronquist 1981, Lupia et al. 1999, Pimm and Joppa 2015, Prance et al. 2000, Thorne 2002*), the presented phylogeny represents between 14% and 8% of the known flowering plants, respectively. In addition, the phylogenetic tree contains 54.6% (8,399) of all currently accepted angiosperm genera, and 94.5% (426) of all families of flowering plants are included, as well as all currently known angiosperm orders. As such, the current angiosperm tree can be regarded as the largest dated angiosperm phylogenetic framework that is generated by combining genuine sequence data and fossil calibration points, and will be useful for large-scale ecological and biogeographical studies. Compared to the species-level based tree of *Zanne et al. (2014)* and its updated version by *Qian and Jin (2016)*, the current phylogeny is larger in size, containing more species (+4,797 species) and genera (+468). However, the phylogeny of *Zanne et al. (2014)* included more families and an equal number of orders. Additionally, *Zanne et al. (2014)*’s updated phylogeny (*Qian and Jin 2016*) also included 1,190 taxa of bryophytes, pteridophytes and gymnosperms, whereas the current phylogeny only contains 5 outgroup gymnosperm species. As a result, when comparing the differences in species number between both angiosperm mega-phylogenies, the current tree contains nearly 20% more flowering plant lineages (+5,987 species).

Age estimation of the large-scale angiosperm tree resulted in a dated phylogeny (*Fig. 1*; *Suppl. material 3*) that largely corresponds to the different recent angiosperm-wide dating analyses (e.g. *Bell et al. 2010, Magallón et al. 2015, Smith et al. 2010, Wikström et al. 2001, Zanne et al. 2014*). Even though small dissimilarities are present concerning the age of the most early diversified angiosperm lineages (see *Table 1*), the overall age of the different families corresponds rather well to what is known from these other studies. Differences in stem node age of large clades such as superasterids, superrosids, eudicots, monocots or magnoliids are probably due to the use of a slightly different and larger set of fossil calibration points, as well as not using tricolpate fossil pollen as maximum-age for eudicots. Compared to the angiosperm phylogeny of *Zanne et al. (2014)*, where time scaling was carried out with 39 fossil calibrations, the current tree contains 56 fossils in total. Although some fossils are the same between both Zanne’s study and ours (e.g. *Pseudosalix handleyi, Fraxinus wilcoxiana, Spirematospermum chandlerae*), several fossils that have been used to optimize the age estimation of the current megaphylogeny are carefully chosen from other dating analyses (*Bell et al. 2010, Magallón et al. 2015, Smith et al. 2010*).

**Figure 1.**
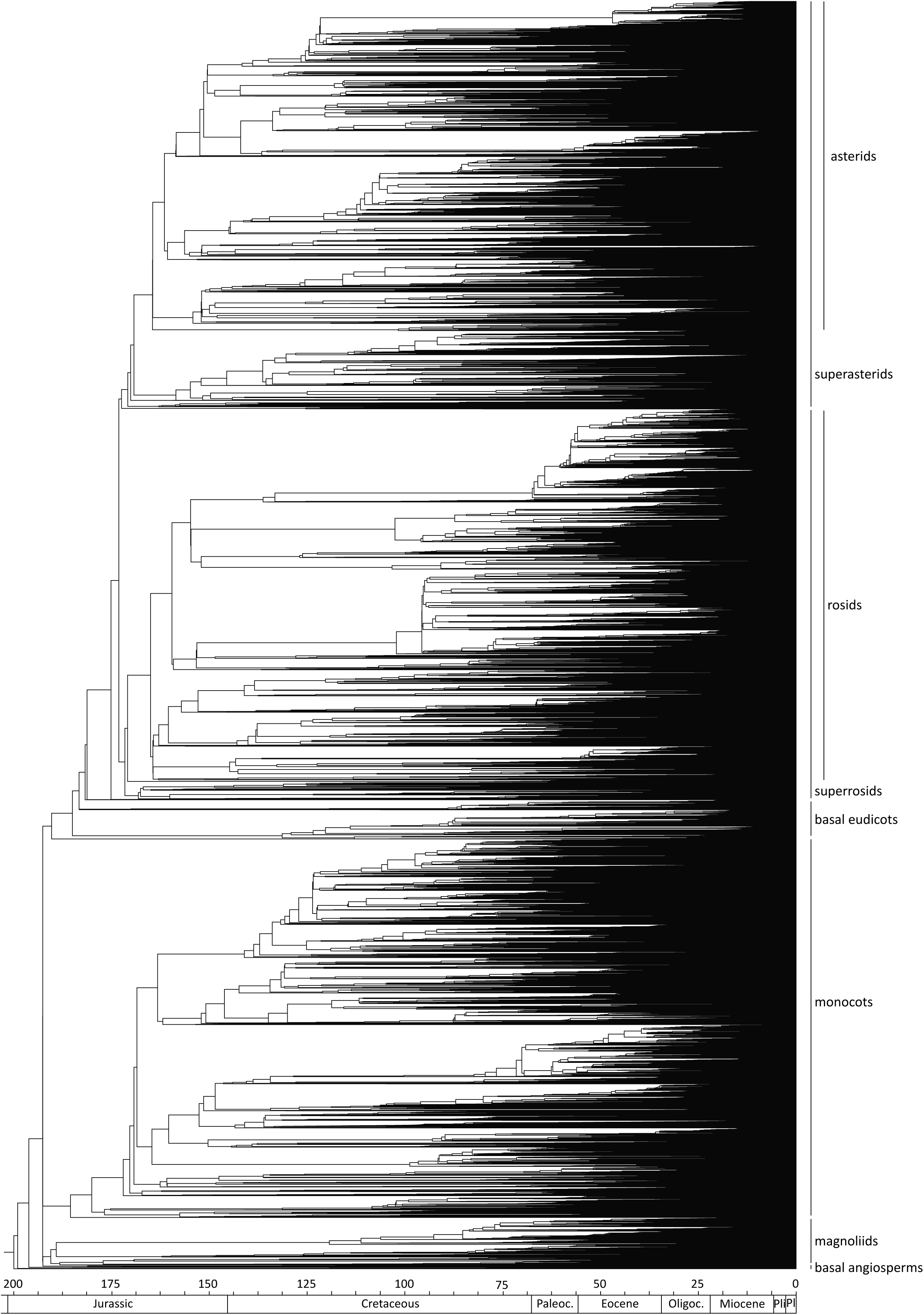
Maximum Likelihood-based angiosperm phylogram based on the combined *rbcL* and *matK* dataset.

Recently, *Qian and Jin (2016)* developed a novel tool (S.PhyloMaker package as implemented in the R environment) to generate artificially enriched species trees based on an updated version of the original angiosperm megaphylogeny of *Zanne et al. (2014)*. According to the study of *Qian and Jin (2016)*, the software package produces phylogenies for every species that one needs to assess in a community ecological environment. S.PhyloMaker grafts species of interest, either as a basal polytomy (regular or Phylomatic/BLADJ approach; *Webb et al. 2008*), or randomly branched within the existing parental clades that are found in the megaphylogeny. Likewise, branch lengths or time-calibrated node splits of newly added taxa are also artificially estimated according to their relative position in the original megaphylogeny. Even though the software package of *Qian and Jin (2016)* provides a good alternative for the lack of descent sampling of angiosperms taxa in megaphylogenies for some ecological studies, not all ecological or evolutionary disciplines that are in need of a phylogenetic framework can rely on this methodology as it is not based on the inclusion of original sequence data. As such, it remains a continuous challenge to increase the size of large-scale angiosperm phylogenies with new species and gene markers to create a reliable platform in which ecological and evolutionary research can be combined with phylogenetics. The current phylogeny is a further step towards an all-encompassing angiosperm phylogeny that can be used to resolve large-scale ecological and evolutionary queries.

## Acknowledgements

This study is part of the HERBAXYLAREDD project (BR/143/A3/HERBAXYLAREDD), funded by the Belgian Belspo-BRAIN program axis 4. This project is supported by Plant.ID, which has received funding from the European Union’s Horizon 2020 research and innovation programme under the Marie Sklodowska-Curie grant agreement N° 765000. This study is also supported by the BRAIN.be BELSPO research program AFRIFORD and by the French Foundation for Research on Biodiversity (FRB) and the Provence-Alpes-Côte d’Azur region (PACA) region through the Centre for Synthesis and Analysis of Biodiversity data (CESAB) programme, as part of the RAINBIO research project (http://rainbio.cesab.org/).

